# Geometric differences in the ribosome exit tunnel impact the escape of small nascent proteins

**DOI:** 10.1101/2022.08.19.504567

**Authors:** Shiqi Yu, Simcha Srebnik, Khanh Dao Duc

**Author notes:** Equal contribution (Simcha Srebnik, Khanh Dao Duc).

## Abstract

The exit tunnel is the sub-compartment of the ribosome that contains the nascent polypeptide chain and as such, is involved in various vital functions, including regulation of translation and protein folding. As the geometry of the tunnel shows important differences across species, we focus on key geometrical features of eukaryote and prokaryote tunnels. We used a simple coarse-grained molecular dynamics model to study the role of the tunnel geometry in the post-translational escape of short proteins (sORF’s), with lengths ranging from 6 to 56 amino acids. We found that the probability of escape for prokaryotes is one for all but the 12-mer chains. Moreover, proteins of this length have an extremely low escape probability in eukaryotes. A detailed examination of the associated single trajectories and energy profiles showed that these variations can be explained by the interplay between the protein configurational space and the confinement effects introduced by the constriction sites of the ribosome exit tunnel. For certain lengths, either one or both of the constriction sites can lead to the trapping of the protein in the “pocket” regions preceding these sites. As the distribution of existing sORF’s indicate some bias in length that is consistent with our findings, we finally suggest that the constraints imposed by the tunnel geometry have impacted the evolution of sORF’s.

## Introduction

The translation of messenger RNA (mRNA) into a protein is a fundamental and complex biological process that is largely mediated by ribosomes. During this process, ribosomes decode the mRNA sequence to assemble the nascent polypeptide chain that transits from the polypeptide transferase center (PTC) to the outer surface of the ribosome, through a sub-compartment known as its exit tunnel. The narrow geometry and composition of the tunnel [1] potentially yield interactions between the nascent chain and the ribosome structure which can significantly impact both translation dynamics [2,3] and protein folding [4–7]. In extreme cases, these interactions can lead to translation arrest as the nascent chain acquires some critical structure (e.g. *α*-helices) within the tunnel [8–10]. Various studies have hence focused on examining the impact of the biophysical properties of the exit tunnel on translation, highlighting key regions where the nascent chain interacts with the tunnel wall [11–13]. Among these key regions, critical interactions are susceptible to take place at the PTC [1, 13], but also downstream at the so-called “constriction site” [12–14] located in the vicinity of the universally conserved ribosomal proteins uL4 and uL22 [15]. A further increase in the tunnel radius at the exit also defines a “vestibular” region, with enough space to accommodate tertiary structure [16, 17]. To analyze these complex dynamics, molecular dynamics simulations were previously used [18] to study how the dynamics of the nascent chain are impacted by various biophysical properties of the tunnel, including its geometry [7, 19–21], electrostatics [5, 22] and hydrophobicity [23]. With the increasing availability of high-resolution structures of the ribosome for various species [24], detailed mappings of the exit tunnel have revealed that these properties can vary across the domains of life [15, 19]. Most notable are the geometric differences in tunnel lengths and diameters, as eukaryotic tunnels are, on average, shorter and substantially narrower than prokaryote ones. [15, 19]. While these differences were recently shown to lead to a faster transit and higher probability of knotting for a specific protein in bacterial ribosomes [19], their implications in some more generalized context are still elusive. In particular, it was suggested that the geometry of the tunnel should especially hinder the escape of smaller polypeptides [20] that can be fully contained in the tunnel (below ∼ 50 amino acids [1]). Interestingly, such polypeptides, also called short open reading frames (sORF’s), have been widely identified with the recent advances in sequencing technology [25], and are involved in some important regulatory functions [26].

In this paper, we examine the extent to which the geometric differences between eukaryotes and prokaryotes can affect the release and escape of sORF’s from the ribosome exit tunnel. To do so, we used a coarse-grained model of the polypeptide chains and simplified tunnel geometries that recapitulate the differences between eukaryotes and prokaryotes. We performed molecular dynamics simulations using this simple representation to study the probability and conditional exit time of the nascent chain, for varying polypeptide lengths. Our comparative approach revealed significant differences between the behavior and escape of proteins in the prokaryotic and eukaryotic tunnels, which we explained upon analyzing in detail the trajectories and energy profiles from our simulations. In conjunction with other recent studies that we discuss below, our study contributes to unraveling the critical and complex role that the tunnel geometry plays in regulating protein translation.

## Material and Methods

### Poplypeptide Model

The polypeptide dynamics were simulated using a coarse-grained model, in which each residue was modeled by a triplet of beads representing the backbone units. For simplicity, all beads were considered to be identical. The short-range intermolecular interactions between non-bonded residues were modeled using the Lennard-Jones (LJ) potential, according to

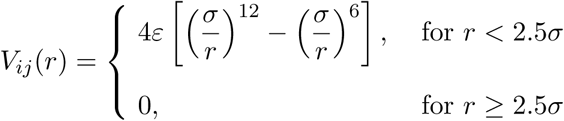

where the LJ parameters, *σ* and *ε* were assigned the values of 3.9 Å and 0.118 kcal/mol, respectively. These values correspond to an alkane unit based on the OPLS-UA force field [27]. Intramolecular interactions were defined using a stiff harmonic bond potential between neighboring residues, given by

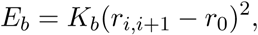

where *r*_*i,i*+1_ is the distance between the centers of two consecutive beads. The bond coefficients of all beads were set to *K*_*b*_ = 270 kJ/mol and *r*_0_ = 1.5 Å. These values are representative of single carbon-carbon bond parameters [27].Angle and dihedral forces were neglected. Polypeptide chains of varying lengths between 6 to 56 amino acids (aa’s) were considered.

### Ribosome exit tunnel model

We considered two simplified tunnel models that are representative of the ribosome exit tunnels of eukaryotes and prokaryotes [15]. The tunnel was modeled as a cylinder of varying radius, aligned along the *z*-axis, with *z* = 0 taken to be the position of the PTC. The radial profile along the eukaryotic tunnel was set to a piecewise constant function with values *r* = [4; 6.5; 4; 6.5; 4; 6.5; 7.5] along the intervals {[0; 10]; [10; 25]; [25; 30]; [30; 50]; [50; 55]; [55; 80]; [80; 90]}, respectively (in Å). These values are the same for the prokaryotic tunnel, except for the radius at the second constriction site (*z* ∈ [50; 55]), where *r* = 5 Å (instead of 4 Å) and at the exit port where the tunnel was 10 Å longer (*z* ∈ [80; 100] instead of [90; 100]). The tunnel is closed at the PTC site and open at the exit port. We included an additional 2 Å for the van der Waals (vdW) radius of a virtual residue embedded in the tunnel wall [28]. The inner surface of the tunnel was treated as a weakly adsorbing rigid bounding wall with short-ranged interactions given by

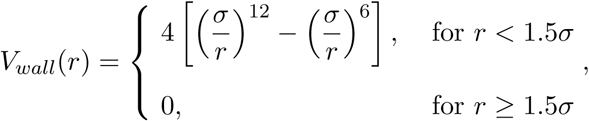

where *r* is the shortest perpendicular distance from an amino acid to the tunnel wall and the LJ parameters were assigned the same values as for the intramolecular LJ potential.

### Simulations of polypeptide dynamics within the tunnel

We used LAMMPS molecular dynamics software [29] to simulate the polypeptide dynamics within the tunnel and its post-translational escape. As the protein translation rate is generally much slower than chain kinetics for the short polymers, the fully translated protein can be considered to be at its equilibrium state for short polypeptide chains [1]. We first equilibrated the linear polypeptide chain in a cylindrical tunnel that is long enough to contain the protein, with a uniform inner surface radius of *r* = 4 Å. The velocity of the protein was initiated at a temperature of 5 a.u., which corresponds to ca. 300 K. Equilibration of the protein required approximately 200,000 timesteps. The protein was then placed in the complex tunnel. The velocity of the protein was reset and the protein was re-equilibrated for at least 200,000 timesteps with its center of mass fixed at *z* = 0 (it generally required much longer to reach equilibrium for proteins longer than 38 residues). We carried out 540 independent runs for polypeptide chains of 20 or more residues in length, and for 240 independent runs for the 6- and 12-long polypeptides. Twenty different configurations were randomly selected as initial states for each tunnel and protein considered (more details are provided in the SI). The conditional escape time of the protein was measured from the moment the last terminal residue was released from the PTC until all the residues exited the tunnel. The maximum simulation time was set to 6,500,000 timesteps. VMD software was used for the visualization of protein structures [30].

### sORF sequences

The sequences used to analyze the sORF length distribution were downloaded from the SmProt database [31] after separately filtering the sORF’s from ribosome profiling sequences in *Homo sapiens* and *Escherichia coli* [25]. Eligible sORF’s were extracted with their sequences, gene name and protein length. We reduced this dataset to distinct genes and lengths of 6 - 40 residues.

## Results

### Simulations of sORF escape from the prokaryotic and eukaryotic ribosome exit tunnels

To examine the impact of the tunnel geometry on the escape of sORF’s, we first simplified its atomic structure with a coarse-grained representation that captures the main differences between prokaryotes and eukaryotes, according to a recent comparative study [15]. As highlighted in Fig. 1a, ribosomal tunnels generally differ between eukaryotes and prokaryotes primarily in the pocket region located between two constriction sites. Due to structural variations of the uL4 protein, the (second) constriction site that is located closer to the exit port is narrower in eukaryotes (on average 4 Å in prokaryotes, and 5 Å in eukaryotes). The prokaryote tunnel is also on average 10 Å longer. Our simplified representation of the tunnel is shown in Fig. 1b, as a cylinderical form of piecewise constant radius. The corresponding radial profiles, plotted in Fig. 1c show in detail the differences between the two tunnel types. We further investigated the role of these two domains on the escape dynamics of small peptides. To do so, we used a coarse-grained model of the polypeptide, where each residue is represented as a triplet of beads with LJ pairwise interactions (see Fig. 1b). In the next sections, we analyze and compare the simulation results for different polypeptide lengths and these two idealized tunnel geometries.

**Figure 1:**
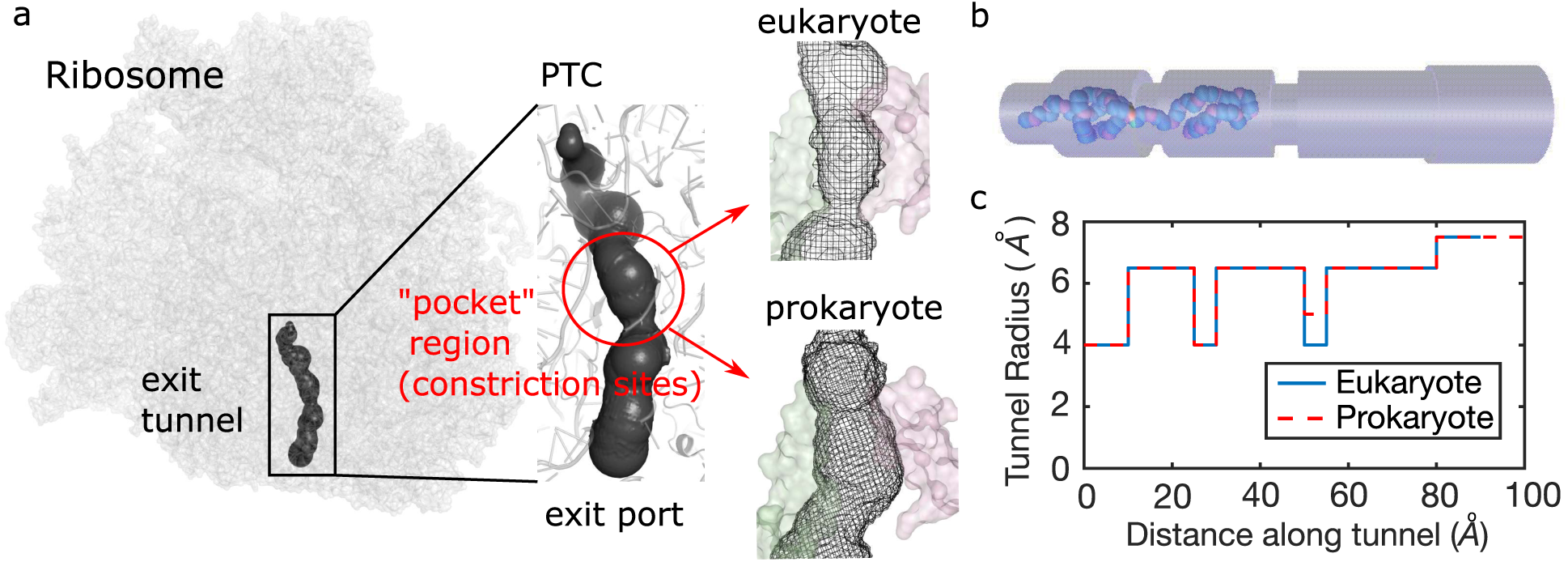
Representations of the exit tunnel for simulating the escape of polypeptides in prokaryote and eukaryote. **(a)** The exit tunnel (black) is a compartment of the ribosome (grey) that extends from the Polypeptide Transferase Center (PTC) to the solvent accessible surface (ribosome structure is taken from [44], with the tunnel extracted as in [2, 15]). The “pocket” region enclosed between the tunnel constrictions are shown in more detail for a eukaryote and a prokaryote (structures taken from [44] and [45]). This region is surrounded by ribosomal proteins uL4 (green) and uL22 (pink), with uL4 forming a narrower constriction site in the eukaryote [15]. **(b)** Snapshot of the coarse-grained nascent chain within a cylindrical domain of piecewise-constant radius representing the eukaryote tunnel (see Material and Methods). **(c)** Radial profile of the prokaryotic (red) and eukaryotic (blue) tunnels used in the simulations (where the origin is set at the PTC).

### Differences in protein escape probability and conditional escape time across tunnel geometries and protein lengths

Our simulations were performed on 6 datasets generated for polypeptide chains with 6, 12, 20, 29, 38 and 56 residues. This range of lengths was chosen since fully extended conformation of these polypeptides can be fully retained in the tunnel. Our simulations focused on the dynamics that follows the release of the polypeptide from the PTC and its escape from the tunnel, that is achieved when all the residues have passed the tunnel exit within the duration of the simulation. Upon recording the statistics of these escape events, we present in Fig. 2 the empirical escape probabilities (*P*) and conditional escape times (*τ*) obtained for the different datasets and tunnel geometries. In the prokaryotic tunnel, all proteins successfully escaped, except for 12-mer polypeptides (with *P* = 0.76). In contrast, we observed that for the eukaryotic tunnel, a significant fraction of polypeptides up to 29 residues in length did not escape, resulting in *P* = 0.72, 0.08, 0.7, and 0.38 for the 6-, 12-, 20- and 29-mers, respectively (Fig. 2a). In particular, the escape of 12-mer polypeptides in eukaryotes was critically hindered, with the vast majority of the chains trapped in the tunnel. Trapping also occurred for the majority of 29-mer polypeptides but to a lesser extent. Despite the eukaryotic tunnel being 10 Å shorter than the prokaryotic tunnel, the conditional mean escape times were significantly longer for polypeptide length ≤ 29 (Fig. 2b). The mean escape time also showed a non-monotonous trend reflecting the inverse of the escape probability. Overall, these results suggest that while variations of tunnel lengths across the domains of life may not significantly influence the polypeptide escape, the geometric variation at the second constriction site produces a strong and non-linear impact on the escape of sORF’s, up to a certain threshold of polypeptide length.

**Figure 2:**
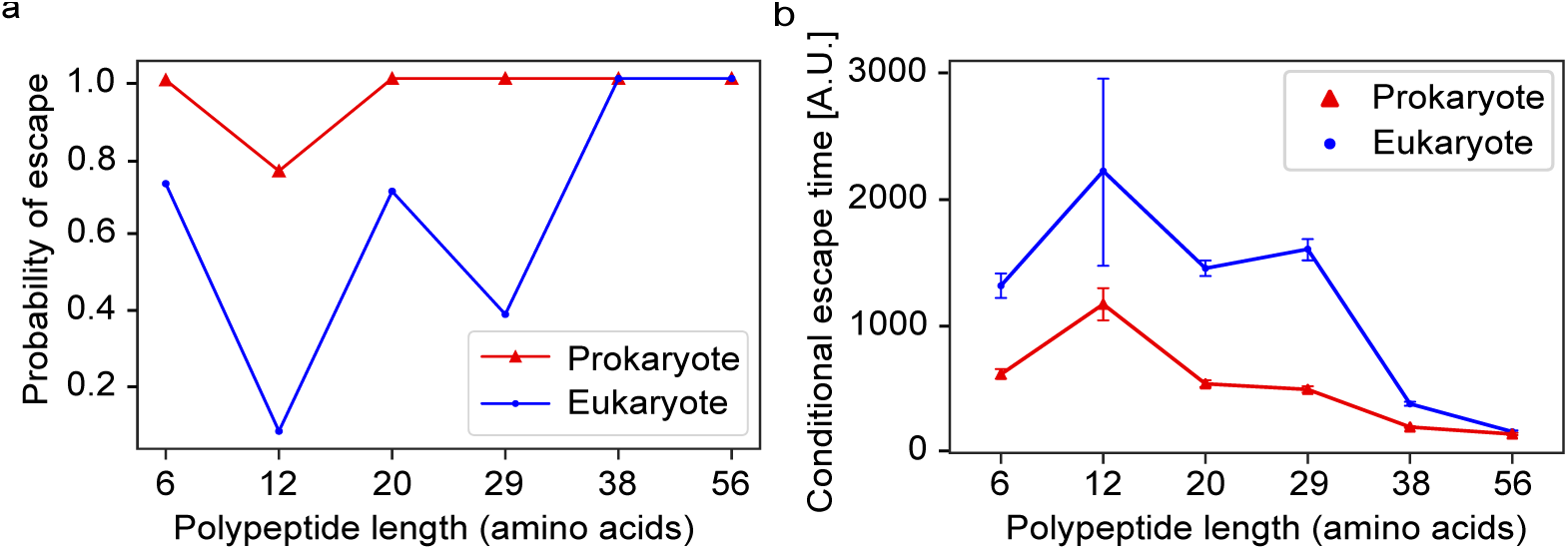
**(a)** Probability of escape and **(b)** conditional escape time (with mean and standard deviation) as a function of polypeptide length for the prokaryotic (in red) and the eukaryotic (in blue) tunnels.

### Polypeptide length-dependent trapping location in the eukaryotic tunnel

To explain the differences in escape probability across protein length, we primarily focused on the eukaryotic tunnel, where trapping was found to be more prominent, and studied the trajectories taken by the polypeptide chain. To do so, we visualized in Fig. 3 the density of the C-terminal trajectories, as well as the distributions of escape time and of the the C-terminus location at the end of the simulation. For the 12-mer proteins with the lowest escape probability, we observed that the C-terminus was susceptible to being stuck in the pocket region between the two constriction sites in 68% of the trajectories (Fig. 3b). In addition, a few trajectories were trapped by the first constriction site (14%). In comparison, the 29-mer proteins, which also had an uncharacteristically low escape probability (Fig. 2a), were surprisingly prone to get indefinitely stuck by the first constriction site rather than the second one. A similar analysis of the density of N-terminal trajectories revealed, however, that the N-terminus of those trajectories that fail to escape are mainly trapped before the second constriction site. Simulation snapshots further revealed that the 29-mer protein was divided by the first constriction site into two similar-sized parts, each completely folded in its respective ‘pocket’, hence reducing the probability of the protein passing through the two narrowest sites simultaneously. We confirmed this result by carrying out a statistical analysis of the time spent by the two termini in different sections of the tunnel. For the cases where proteins were unable to escape, the two termini spent around 64% of the total simulation time simultaneously stuck in the different pockets, suggesting that each terminus was restricted by a different constriction site. Therefore, when the second constriction site was less narrow, as in prokaryotic tunnel, the 29-mer proteins were successfully drawn out of the tunnel by their N-terminus, resulting in a higher escape probability and a lower escape time (Fig. 2).

**Figure 3:**
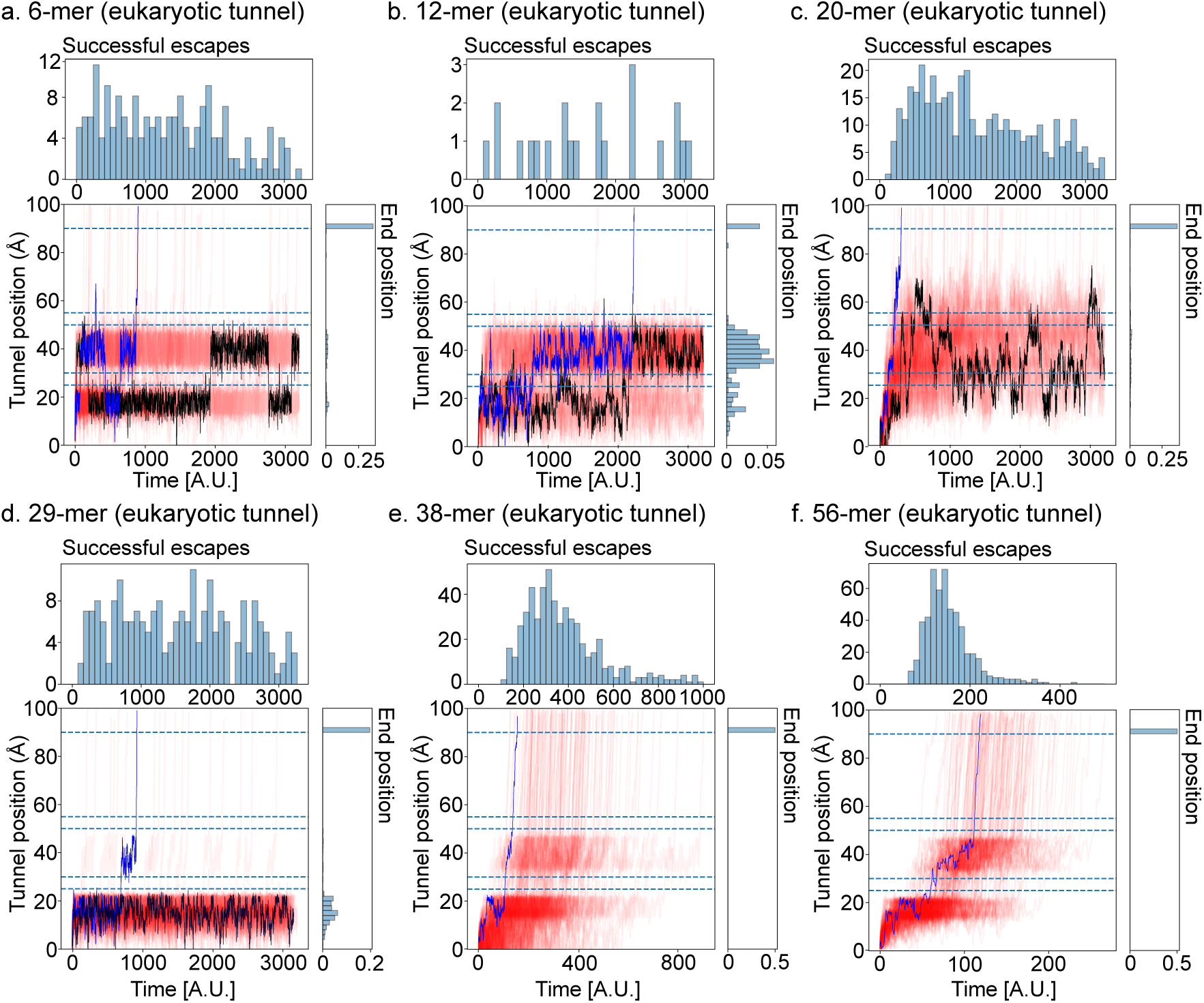
C-terminus trajectories in the eukaryotic tunnel. For each dataset (polypeptide length 6 to 56 **(a-f)**), the positions of C-terminus along the tunnel over time for all simulated proteins of a given length are plotted in red. We superimposed representative trajectories of the polypeptide escaping (in blue) or being trapped (in black). The dashed lines at *z* = 25, 30, 50, and 55 Å indicate the beginning and end location of the two constriction sites. The dashed lines at *z* = 90 Å indicate the location of the tunnel exit. Upper histograms show the conditional exit time distribution for each dataset, while histograms on the right show the distribution of C-terminus final position.

For the 6-mer and 20-mer proteins (which showed comparable escape probability), we found that 54% and 73%, respectively, of C-terminus trajectories float back and forth within the tunnel for long periods of time (Fig. 3a, c). These proteins readily pass through the first constriction site while a small number is obstructed by the second one. Overall, shorter proteins (*<*38 aa) were thus significantly affected by the second constriction sitein the eukaryotic tunnel.

Finally, the trajectories of the longest proteins (*>*=38 aa) showed that the C-terminus successively transits across the PTC to the pocket and the exit regions (Fig. 3e, f). The corresponding N-terminal trajectories notably showed that the initial positions of the N-terminus (once the polypeptide was released from the PTC) were beyond the first constriction site for the 38-mer proteins and were even well beyond the second constriction for the 56-mer proteins. Since the N-terminus was rarely blocked by the constriction sites for these longer proteins, it was entropically driven towards to the tunnel exit, thus sequentially drawing the remaining segments, leading to rapid escape without being trapped.

### Energy profiles in the tunnel

Our results suggest that the primary hindrance of the escape of small proteins is the second constriction site present in eukaryotes. The translocation can be roughly estimated in terms of the N-terminus position since it was found to be closely correlated with the escape probability. However, it is still unclear why the 12-mer protein has the lowest escape probability in both tunnels. To address this, we focused here on the 12-, 20-, 29-mer proteins, and evaluated from our simulations the potential energy of the nascent polypeptide and the contribution of intramolecular interactions at different locations along the tunnel. In Fig. 4, we show the resulting plots of the average intramolecular interaction energy of the protein as a function of the N-terminus position, leading to distinct energy landscapes. The energy was calculated every 10,000 timesteps by MD simulations and the average was over all trajectories for each dataset.

**Figure 4:**
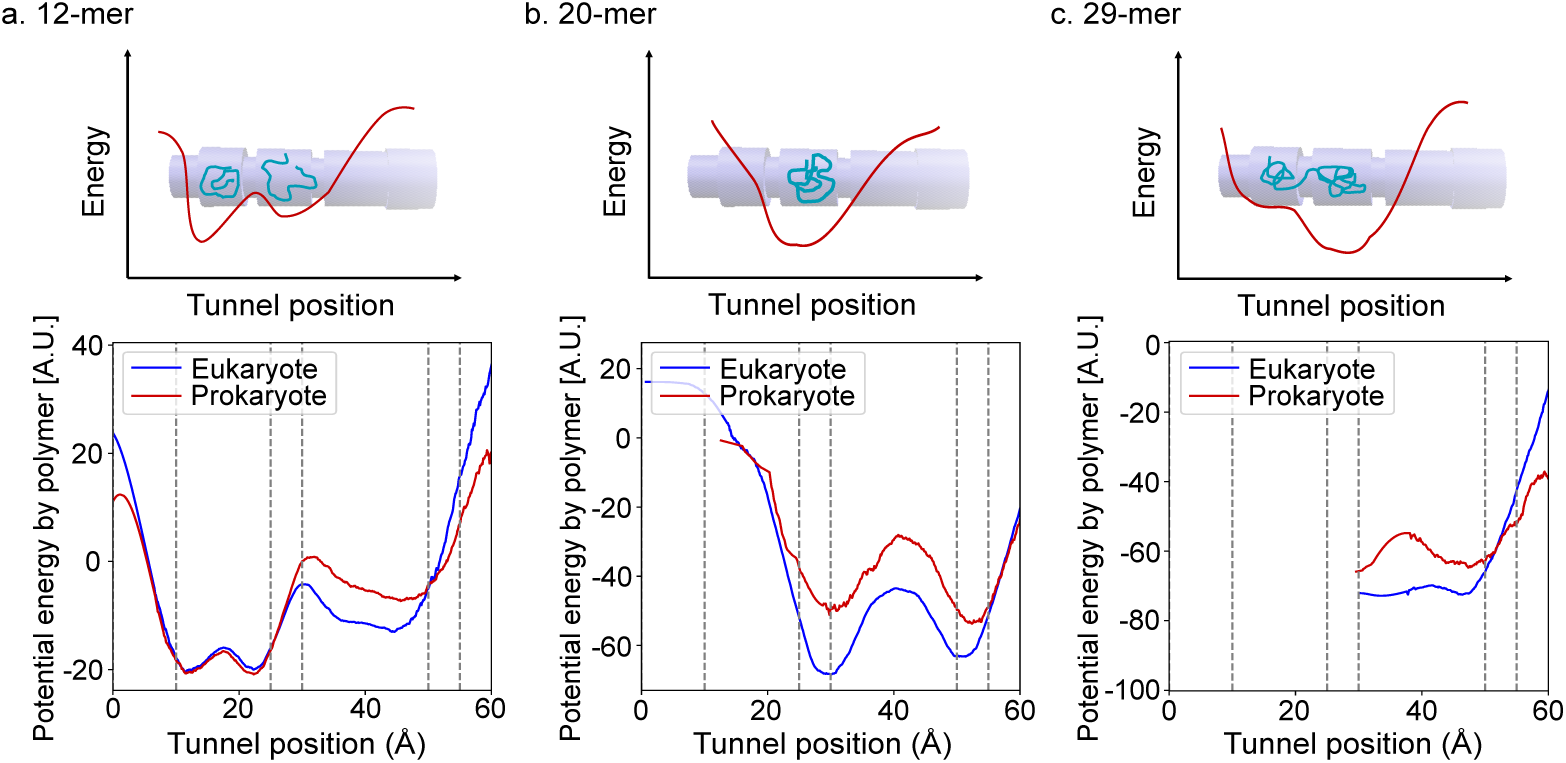
Intramolecular energy of the proteins. The average potential energy of proteins in the eukaryotic tunnel for **(a)** 12-, **(b)** 20-, and **(c)** 29-mer chains as a function of the N-terminal position. The dashed lines represent the start of the first pocket and the positions of the constriction sites. A Savitzky-Golay filter was applied to reduce the noise. The upper plots show the general trend of the free energy of the protein in the tunnel and a schematic of intermediate protein configurations.

For the 12-mer proteins (Fig. 4a), the polypeptide experienced lower energy in the first smaller pocket immediately following the PTC. This small space is just sufficient to accomodate the entire protein, leading to significant intramolecular interactions (see Fig. 4a), as well as favorable interactions with the wall. The low energetic state in this pocket is counteracted by the entropic penalty of confinement and the strong entropic drive towards the exit introduced by the closure of the tunnel at the PTC end [32]. The larger second pocket (between the two constriction sites) allows for less compact conformations for the 12-mer protein, leading to an overall higher intramolecular energy. While the drive to escape from the second pocket is substantially lower for the 12-mer because of its higher entropy [33], both the first and second constrictions present a significant thermodynamic hindrance, slowing down its escape. The wider second constriction of the prokaryotic tunnel is sufficient for the protein to easily overcome this energy barrier in favour of the higher entropy of the free chain [33]. In contrast with the 12-mer, the first pocket is too small to fully accommodate the 20-mer proteins, resulting in unfavourable intramolecular interactions (Fig. 4b). This low entropy and high energy state readily drives these proteins towards prompt escape, facilitating their passage through the confinements. Accounting for the preferred vdW interaction radius of 1.12 *σ*, the first and second pockets can ideally accommodate between ca. 15 and 20 monomers, respectively. The second constriction therefore presents a higher thermodynamic barrier for the 20-mer to escape. For the 29-mer proteins (Fig. 4c), the initial position of the N-terminus was found to be already beyond the first narrow region, so that a large energy well is seen between the two constriction sites. The 29-mer proteins comfortably occupied the two pockets simultaneously. Escape requires overcoming this low energy state and the N-terminus passing through the second constriction site. The initial position of the N-terminus of even longer proteins was already beyond the second constriction, so that no entropic barriers were present for the proteins’ escape.

### sORF sequence analysis

We finally studied the potential evolutionary implications of the polypeptide trapping in the tunnel by analyzing the size distribution of sORF’s found in both eukaryotes and prokaryotes. We focused on *H. sapiens* and *E. coli*, as their sORF sequences have been recently identified and annotated using ribosome profiling [31]. After selecting gene sequences of lengths less than 40 codons, we obtained a histogram of sORF’s lengths for both species, plotted in Fig. 5. In both distributions, we found a significant dip in the number of sORF’s for proteins around 10 amino acids in length, which matches the length at which the escape probability was critically low in our simulations (Fig. 2a, b). More precisely, we found 696 sORF sequences of protein length comprised between 8 and 13 amino acids in Human, compared with 1203, 1069, 1036 and 867 sequences of length comprised between 14-19, 20-25, 26-31 and 32-37 amino acids, respectively. While the number of sORF’s was one order of magnitude smaller in *E. Coli* (5784 sequences in Fig. 5a, compared with 365 in Fig. 5b), we found the same trend in the histogram as in the human sORF’s, with 34 sequences found between 8 and 13 amino acids, compared with 62, 78, 66, 68 in the remaining parts of the histogram. Upon running a statistical *χ*^2^ test of goodness-of-fit with the uniform distribution, we found that one cannot reject the null hypothesis that sequences are uniformly distributed (*χ*^2^ = 2, with p-value = 0.56 *>* 0.05) between length 15-40 in *E. coli*. However, the same hypothesis was rejected when adding observations from length 8-14 (*χ*^2^ = 17.9 with p-value = 0.001 *<* 0.05), suggesting some possible negative selective pressure against sORF’s at the specific size where the geometry of the exit tunnel hinders the escape of the nascent polypeptide. Performing the same *χ*^2^ tests in human led to reject the null hypothesis in both cases (with a smaller p-value found when including all sequences from length 8 to 40). In this case we also observe in the 15-40 amino acids region a downwards trend in the number of sORF’s as the polypeptide length increases (Fig. 5a), which cannot be simply explained by the variations of escape probability that we obtained from our simulations.

**Figure 5:**
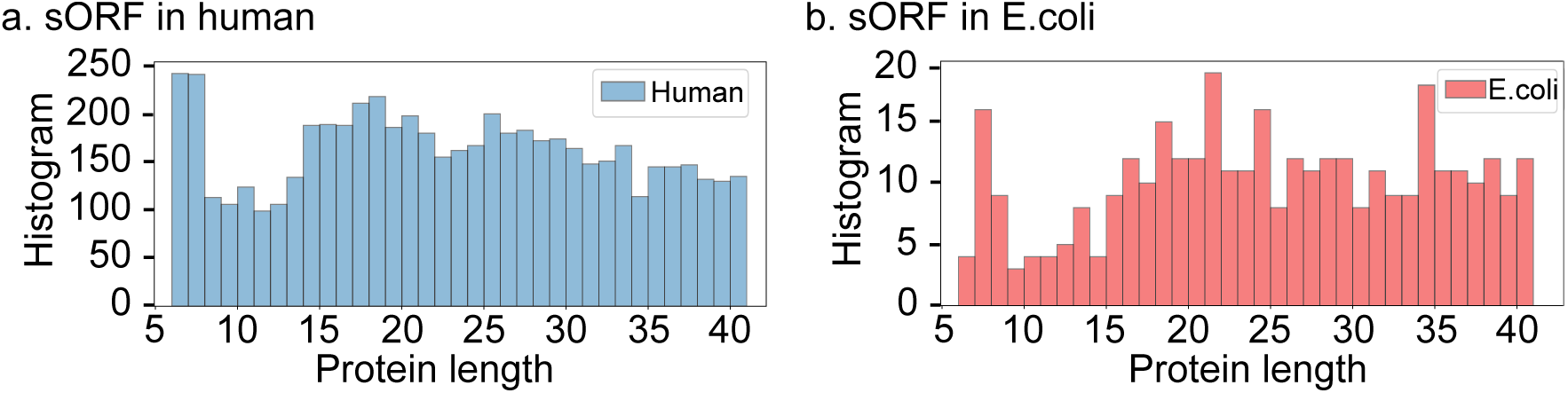
Length distribution of existing sORF’s with less than 40 amino acids, in **(a)** Human and **(b)** *E. coli*. Data was collected from SmProt database [31] (see Material and Methods.

## Discussion

We used coarse-grained molecular dynamics to study the escape process of sORF’s from the ribosome exit tunnel. Our simulations revealed that these short polypeptide chains can significantly get trapped during this process, with a probability that not only depends on the polypeptide length, but also differs between eukaryotes and prokaryotes. Analyzing the trajectories of the polypeptides and energy profiles associated with the tunnel interactions allowed us to explain these variations, and how the trapping occurs in specific regions of the tunnel, where high energy barriers limit the entropic drive for the protein to move downstream. In particular, our study highlights the role played by the constriction sites and the “pocket regions” they enclose, as escape is more difficult when the size of the partially folded conformation is comparable to the spatial volume of these pockets. The entropy associated with excluded volume drives towards translocation of the protein through the constrictions. This drive increases logarithmically with chain length, but the escape time decreases exponentially with the increasing size of the constriction [33].

While various studies have also used MD simulations over the past few years to study the polypeptide escape and the impact of the tunnel properties (including [19, 22, 28, 34]), our study is, to our knowledge, the first to comprehensively study how trapping occurs post-translationally in the exit tunnel at the scale of small sORF’s. For larger proteins, Bui and Hoang compared various single-domain proteins of length comprised between 37 and 99 residues [20], and observed that these proteins can successfully escape from the tunnel, in agreement with our results for polypeptides of length ≥ 38. Interestingly, they also simulated a synthetic 29-residue zinc-finger domain, and found it to be severely and deeply trapped within the tunnel. Our study confirms that such a trapping is more prone to happen at this size, and provides a more complex picture that shows that the extent of this phenomenon varies according to the size of the organism considered. The complex variation in protein translocation based on its size and the organism suggests some potential functional implications for the translation of sORF’s.

Various studies also showed over the past few years how the constriction site can impacts multiple cellular processes, including translation dynamics, protein quality control, antibiotic resistance or co-translational folding [5, 35–37]. Using MD simulations to model the dynamics of a bacterial nascent chain in six rigid all-atom tunnel structures from prokaryotes, archaea and eukaryotes, Chwastyk and Cieplak found that the narrower second constriction site in bacteria enhances folding and knotting stronger than the other ribosomes, while trapping and arrest are more prone in eukaryotic and archaeal tunnels [19]. Aside from the fact that we focused on post-translational escape of sORF’s (while they examined co-translational escape), our study also differs in using a simplified representation of the tunnel as a straight cylinder, similar to some other recent theoretical studies of the exit tunnel [38, 39]. While our representation still allows to capture the main differences found between the eukaryotic and prokaryotic tunnels [15], it would be interesting to extend our study to some less coarse-grained and more realistic tunnels, and possibly study the variability of the polypeptide escape across species of the same domain.

In addition to the geometric aspects of the tunnel, which are the main focus of this study, it would also be interesting to take into account some other biophysical properties of the tunnel and nascent chain [22, 34]. We note that our protein datasets included various charge distributions, so that the polypeptide chain also underwent electrostatic pairwise interactions between monomers. However, we did not observe a discernible effect on protein escape based on the amount and distribution of charge. This suggests that the electrostatic environment created by the ribosome is the main factor modulating the protein escape when it contains charged residues. MD simulations that included electrostatic tunnel-peptide interactions indeed showed that nascent proteins with abundant negatively charged residues near their C-terminus can eject faster (with an opposite effect for positively charged residues) [22]. Using high throughput sequencing data, we also previously reported that gene-specific elongation rates are modulated at early stage of translation (first 10 amino acids) by the presence of charged and polar residues in the polypeptide chain, suggesting that the electrostatic potential and hydropathy of the tunnel can impact the transit of the protein [2]. The same study also showed that the observed frequencies of charged residues in protein sequences are also biased, suggesting some evolutionary pressure for optimizing charge distributions to facilitate elongation and escape through the tunnel. Similarly, we suggest here that the constraints imposed on the escape process by the exit tunnel could result in some selective pressure over the sORF’s size (see Fig. 5). As the variations in escape probability that we found are only partially able to explain the size bias found in human data, it would be interesting to more thoroughly investigate the composition of sORF’s sequences, to fully understand the impact of the biophysical constraints imposed by the tunnel on their evolution. Finally, we suggest some analytical and experimental complementary approaches to the present study, to confirm and extend our findings. On the theoretical side, we note that a thorough study of the interplay between kinetics and thermodynamics around the constriction sites is still required, beyond the results obtained from Fig. 4, to unravel the general relation between the geometry of the tunnel and the escape probability of the nascent chain. Since the energy profiles across the two domains of life is quite similar, the differences in escape times for a given chain length were qualitatively explained based on polymer translocation scaling theory [33]. In principle, one can also analytically formulate the escape process of the nascent chain as a diffusion process of a polymer in a confined domain, and study the first passage time problem associated with its translocation [40]. Methods developed to solve this problem include the Fick-Jacobs approximation, that allows to reduce the PDE associated with the diffusion process over the longitudinal direction of a cylinder [40, 41]. It would thus be interesting to apply such methods in the context of the ribosome exit tunnel, and for various radial and electrostatic profiles deriving from existing structures. Experimentally, optical tweezers have proven useful to measure study ribosome stalling, and evaluate the mechanical forces associated with the tunnel [5, 42], but such methods would be challenging to apply for sORF’s that are deeply buried within the tunnel. Alternatively, some further validation of our results could be done by using cell-free protein synthesis systems, available for both eukaryotes and prokaryotes [7, 43], and testing the outcomes of *in vitro* translation for various mRNA sequences, corresponding to the size of the proteins in our datasets.

## Acknowledgments

This research is funded by NSERC Discovery Grants (PG 22R3468 and PG 11R02115) and a UBC STAIR grant. Computational resources and services were provided by Compute Canada (www.computecanada.ca). We thank A. Kushner for his help.

The authors declare no competing interests.

